# Magnetic Resonance Spectroscopy Spectral Registration Using Deep Learning

**DOI:** 10.1101/2022.08.31.506120

**Authors:** David J. Ma, Yanting Yang, Natalia Harguindeguy, Ye Tian, Jia Guo

## Abstract

A novel convolutional neural network based spectral registration (CNN-SR) approach is introduced to achieve efficient and accurate simultaneous frequency-and-phase correction (FPC) of single-voxel MEGA-PRESS magnetic resonance spectroscopy (MRS) data. For this approach, one neural network was trained and validated using a published simulated and *in vivo* MEGA-PRESS MRS dataset with a wide-range of artificial frequency (0-20 Hz) and phase (0-90°) offsets applied. The proposed CNN-SR approach was subsequently tested and compared to the sequential FPC deep learning approaches and demonstrated more effective and accurate performance. Furthermore, a large random Gaussian signal-to-noise ratio (SNR 20 and SNR 2.5) and line broadening (0-20 ms) was introduced to the original simulated dataset to investigate our model performance compared to the other deep learning models. The testing showed that CNN-SR was a more accurate quantification tool and resulted in a lower SNR when compared with the other deep learning methods, due to having smaller mean absolute errors in both frequency and phase offset predictions. For Off spectra, the CNN-SR model was capable of correcting frequency offsets with 0.014 ± 0.010 Hz and phase offsets with 0.104 ± 0.076° absolute errors on average for unseen simulated data with SNR 20 and correcting frequency offsets with 0.678 ± 0.883 Hz and phase offsets with 2.367 ± 2.616° absolute errors on average at very low SNR (2.5) and line broadening (0-20 ms) introduced. In addition, we tested the simulated dataset with additional SNR and line broadening on a more refined model (CNN-SR+) where the pre-trained CNN-SR was further optimized by minimizing the difference between individual spectra and a common template with unsupervised learning. The performance on Off spectra was improved to 0.058 ± 0.050 Hz for correcting frequency offsets and to 0.416 ± 0.317° for correcting phase offsets. We further evaluated the ability of our model to process the published Big GABA *in vivo* dataset and CNN-SR+ achieved the best performance. Moreover, additional frequency and phase offsets (i.e., small, moderate, large) were applied to the *in vivo* dataset, and CNN-SR+ also demonstrated better performance for FPC when compared to the other deep learning models. These results indicate the utility of using deep learning for spectral registration and demonstrate the application of unsupervised learning in further improving the model to achieve state-of-the-art performance.

## 1. Introduction

Spectral Registration is a widely used technique to correct frequency and phase offsets. This algorithm is designed to align individual transients to a spectra template using a least square fitting method by maximizing the cross-correlation. This technique is widely implemented in spectral editing software and applied on Magnetic Resonance Spectroscopy (MRS) data [1, 2, 3]. MR Spectroscopy (MRS) is an analytical tool used to quantify metabolic chemical changes in human and animal brains, which can provide crucial information on brain health. However, due to MR being highly sensitive to scanner variabilities such as frequency drift and subject motion, frequency and phase shifts may arise, affecting data analysis. Thus, frequency-and-phase correction (FPC) is an essential step for accurate spectral registration and metabolite quantification. For instance, the metabolite GABA is the primary inhibitory neurotransmitter in the human brain, but its concentration is very difficult to quantify due to the overlapping metabolite Creatine (Cr) being present in much greater concentrations. Among a wide range of techniques to assess GABA *in vivo*, MEGA-PRESS is now the most widely used MRS technique [4, 5, 6]. MEGA-PRESS is a J-difference editing (JDE) pulse sequence that separates overlapping metabolites from each other. However, a major limitation in JDE pulse sequences is that they hugely rely on the subtraction of spectral edited “On Spectra” and non-edited “Off Spectra” to reveal the edited resonance in the “Diff Spectra”. As a result of the overlapping resonances being an order of magnitude larger in intensity than the GABA resonance, small changes in scanner frequency and spectral phase will lead to incomplete subtraction in the edited spectrum. The standard approach in GABA editing is to apply frequency and phase drift correction of individual frequency domain transients [4, 7] by fitting the Cr signal at 3 ppm. The major limitation of the Cr fitting-based correction method, however, is that it relies strongly on sufficient SNR of the Cr signal in the spectrum. To overcome this limitation, the previously mentioned spectral registration (SR) approaches were proposed that can accurately align single transients in the time domain [2, 3] or frequency domain [1]. However, the correction accuracy largely depends on the overall spectral SNR where low SNR (i.e. 2.5) will deteriorate the performance as the signal will be dominated by noise. Furthermore, due to increasingly demanding medical needs, it is crucial to develop a more robust, fast and high registration accuracy technique.

To address incomplete spectra subtraction and low registration efficiency, deep learning, a popular technique used to address complex computational challenges, has been an effective and successful image processing tool adopted in medical image registration [8, 9]. This learning-based registration method optimizes a global functional for a dataset during training, thereby limiting time-consumption and computationally expensive per-image optimization during inference. A multilayer perceptron (MLP) model [10] and a convolutional neural network (CNN) model [11] have been recently applied to single-transient sequential FPC for edited MRS. Both of these models (MLP-FPC and CNN-FPC) demonstrate the great potential of applying deep learning in MRS data preprocessing by pre-training models with simulated datasets with wide ranges of frequency and phase offsets. Although both these models yield well-predicted results, the utility in spectral registration is limited due to the models’ need to be separately trained for frequency and phase offset prediction, and separately used to perform FPC. Thus, a more efficient network that can mimic the nature of performing simultaneous FPC similarly to the spectral editing techniques could be considered to more accurately perform FPC of the given data. A CNN spectral registration model (CNN-SR) is considered to correct frequency and phase offset at the same time while comprising the CNN properties of exploiting spatial and temporal invariance in recognition of features such as the overall shape of the signal and its peaks.

In this study, we aim to investigate the feasibility and utility of CNNs for spectral registration of single voxel MEGA-PRESS MRS data. We implemented a CNN spectral registration (CNN-SR) model that performs simultaneous FPC. Our proposed CNN-SR approach, was tested on a published simulated dataset and an *in vivo* dataset against the benchmark neural network approach using MLP and CNN [10, 11]. The model achieved superior performance when compared to MLP-FPC and CNN-FPC. The model performance with additional noise and line broadening of SNR 2.5 and 0-20 ms, respectively, was tested to further demonstrate the utility of CNN-SR when dealing with a more distorted spectra. In addition, a more real-life scenario was simulated by using *in vivo* MRS data with different magnitude of additional offsets (none, small, moderate, large) to further demonstrate the utility of our model to accurately predict the spectral frequency and phase offsets. We devised an unsupervised learning spectral registration technique and applied it to the CNN-SR+ on the *in vivo* data. The CNN-SR+ model performed better than MLP-SR, CNN-SR and had similar performances with the published numerical method model-based SR (mSR) [2].

## 2. Methods

### 2.1. Datasets

#### 2.1.1. Simulated Datasets

The main challenge for deep learning is to determine the inputs and ground truth for networking training in order to achieve a specific performance goal. Since there is no ground truth of frequency and phase offsets for the *in vivo* dataset, in this work, the MEGA-PRESS training, validation, and test transients were simulated using the FID-A toolbox (version 1.2) in Matlab, with the same parameters as described in the previous work [10, 11]. The training set was allocated 32,000 OFF+ON spectra, and 4,000 for both validation set and test set. We also tested our model using datasets with added random Gaussian noise at SNR 20 and further challenged the model with lower SNR 2.5 and Line Broadening (0 - 20 ms). The SNR values were computed by the ratio of the Cr peak signal relative to the noise standard deviation.

#### 2.1.2. In vivo Datasets

*In vivo* data was retrieved from the publicly available Big GABA repository [12]. All 101 MEGA-edited datasets from nine sites with Philips scanners were collected in total where each dataset contained 320 transients OFF+ON. The models were also evaluated on this Philips *in vivo* dataset with additional offsets (small, medium, large).

### 2.2. Network architecture

Both supervised and unsupervised learning were incorporated in our proposed CNN-SR model on the simulation dataset, which was further fine tuned using unsupervised learning on the *in vivo* dataset. During supervised training, both supervised loss and and unsupervised loss were implemented to optimize the network parameters (Figure 1A). Based on the CNN-SR model, CNN-SR+ model only used unsupervised loss which was represented as solid line in Figure 1A.

**Figure 1:**
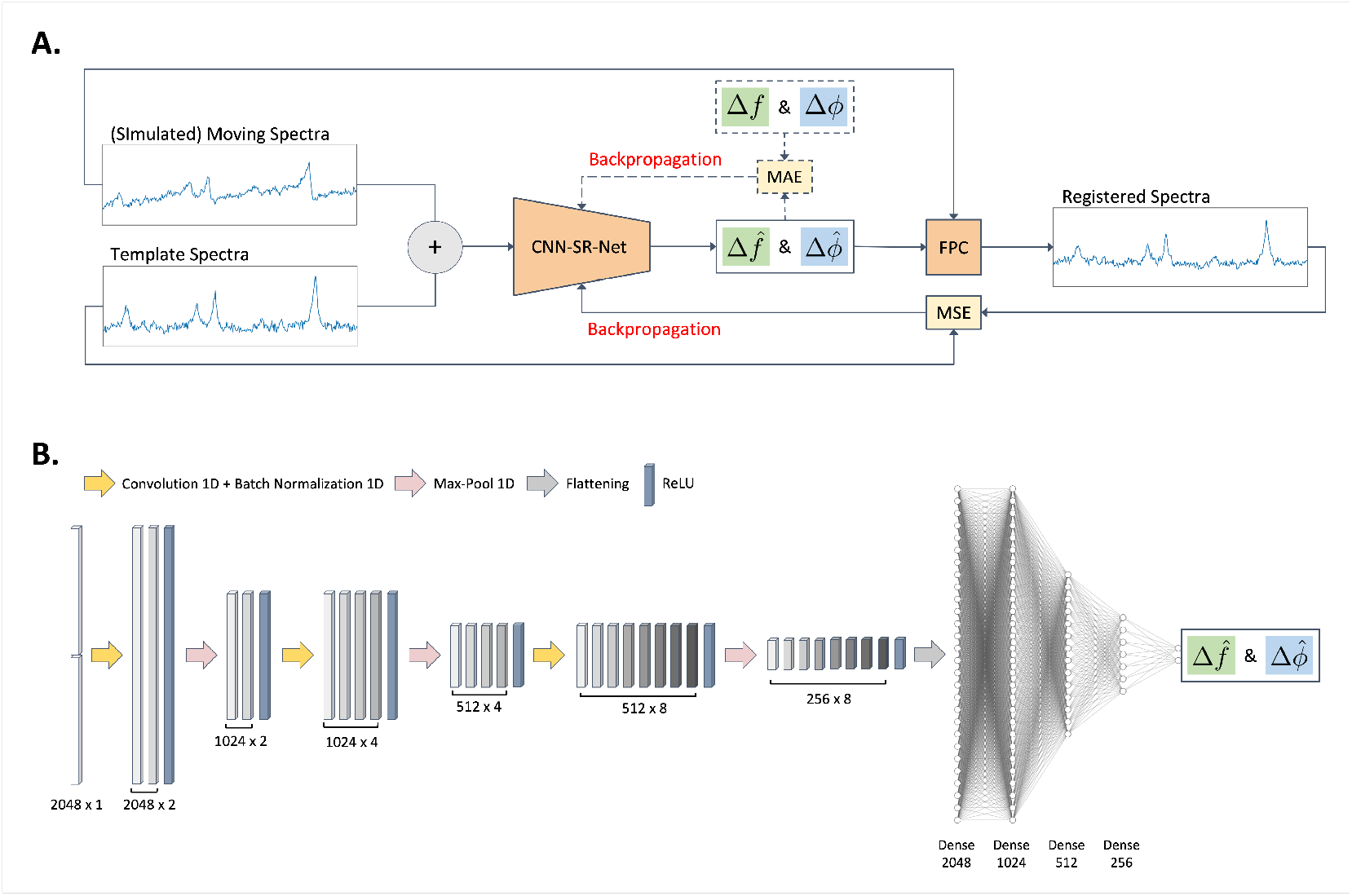
The pipeline for assessment, sample output and network structure of our model. (A) Flow chart of computation to determine the registered spectra with details of the input and output from the network architecture. (B) The network architecture of the CNN-SR model. Both the frequency and phase offsets were predicted with the proposed model where the input is the concatenation of the moving spectra and template spectra. The network architecture was composed of 3 hidden 1D convolutional layers, 3 1D max-pooling layers, and 4 fully connected layers. The convolutional layer consisted of kernels with a size of 3, and the max-pooling layer had a pool size of 2 with a stride of 2. Furthermore, three fully-connected layers (FC) with 1024, 512 and 256 nodes respectively followed by a final fully-connected linear output layer of 2 node, were implemented. All hidden layers were each followed by a rectified linear unit (ReLU) activation function and the output fully connected layer by a linear activation function that generated the predicted offset. Simulated spectra manipulated from FID-A with artificially generated frequency or phase offsets were used as training data for the network. To compare different models, each network was trained through 300 epochs with early stopping implemented when 40 consecutive epochs did not improve the lowest validation loss.

The structure of the network (Figure 1B) was a sequential network which took moving spectra and template spectra as inputs and predicted frequency and phase offsets at the same time. Both moving spectra and template spectra were processed to have length of 1024 and were concatenated to form a single 2048 input array. The network started with three successive layers, each consisted of an one-dimensional convolutional layer followed by a one-dimensional max-pooling layer. The convolutional layer consisted of 4 kernels with a size of 3, and the max-pooling layer has a pool size of 2 with a stride of 2. Furthermore, three fully-connected layers (FC) with 1024, 512 and 256 nodes were used and a final fully-connected linear output layer of 2 node was designed. Each hidden layer was followed by a rectified linear unit (ReLU) activation function to introduce non-linearity. An Adam optimizer [13] was used to train the neural network with a 0.0001 learning rate. The output from the network were the predicted offsets of frequency and phase. Each model was trained for 300 epochs with a batch size of 32, and the mean absolute error was used as the loss function.

### 2.3. Network testing

On the scale of -20 to 20 Hz and *−*90° to 90°, uniformly distributed artificial offsets were first added to simulated spectra to generate input moving spectra, consisting of a frequency drift and a phase drift. Different level of Gaussian distributed noise and Line Broadening was added to the moving spectra prior inputting into the network. First, we applied a Fast Fourier transform to the uncorrected moving spectra and normalized them to the maximum signal in the spectrum. The peripheral 1024 samples were then cropped off, and the central 1024 samples were selected and the absolute value was taken to feed the network. Next, we applied the same normalization and cropping processes to the template spectra and concatenated them into array of length 2048. The network predicted the frequency offset (Δ*f*) and the phase offset (Δ*ϕ*), which were applied to the moving spectra using FPC to generate the registered spectra.

#### 2.3.1. Evaluation and comparison using in vivo dataset

The MEGA-edited datasets were used as the test set of the CNN-SR network. For a first comparison to the performance of our CNN model, a published model-based SR (mSR) [2], a non-deep learning approach, was used to perform FPC in the time domain. mSR uses a noise-free model as the template instead of the median transient of the dataset. Noise-free ON and OFF FID models were created in Osprey (version 1.0.0), an open-source MatLab toolbox, following peer-reviewed preprocessing recommendations [12]. Our CNN-SR model was also compared to a benchmark neural network using MLP containing 3 FC layers (1024, 512, 1 node(s)) [10] and a CNN containing two convolutional blocks (Convolutional layer with 4 kernels of size 3 + Max pooling layer with downsampling size 2 and stride 2) and 3 FC layers (1024, 512, 1 node(s)) [11]. In both of these networks, each hidden FC layer was followed by a ReLU activation function, and a linear activation function followed the output layer.

To examine the network in a more extreme environment, additional series of artificial offsets were added to the *in vivo* data. There were 3 different kinds of additionally added offsets: 1. 0 ≤ |Δ*f*| ≤ 5 *Hz* and 0° ≤ |Δ*ϕ*| ≤ 20°; 2. 5 ≤ |Δ*f*| ≤ 10 *Hz* and 20° ≤ |Δ*ϕ*| ≤ 45°; 3. 10 ≤ |Δ*f*| ≤ 20 *Hz* and 45° ≤ |Δ*ϕ*| ≤ 90°. All additional offsets were sampled from a uniform distribution and added as random pairs of frequency and phase to each transient.

#### 2.3.2. Hardware and software description

This research was conducted with an Intel (R) Xeon (R) CPU E5-2650 v4 @ 2.20 GHz processor and an NVIDIA GeForce RTX 2080 Ti GPU with a memory of 11 GB.

### 2.4. Performance measurement

In the simulated dataset, the artificial offsets were set as the ground truth, and the mean absolute error between the ground truth and predicted value was used as the criteria to measure the network’s performance. Moreover, we calculated and plotted the difference value between the true spectra and the corrected spectra using mSR, MLP-FPC, CNN-FPC and CNN-SR. A Q score [10] was used to determine the performance strengths of each different methods, and it is defined as 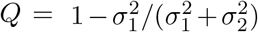, where *σ*^2^ is the variance of the choline subtracted artifact in the average difference spectrum. If the Q score is greater than 0.5, it indicates that the first method performs better than the second method and vice versa.

## 3. Results

### 3.1. Model-performance Evaluation and Spectra Analysis for the Simulated Datasets

The results of the MLP-based approach and CNN-based approaches on the simulated test dataset with lower SNR of 20 and SNR of 2.5 with line broadening are illustrated in Figure 2 and Figure 3. In each subfigure, the frequency offset errors are plotted against their corresponding correct values, the phase offset errors are plotted against their corresponding correct values, the model-corrected difference spectrum and the difference spectrum corrected by the true offsets are plotted together, and the residues between the difference spectra are shown. The comparison of the errors for FPC of the MLP-based approach and the CNN-based approach of the On, Off and Diff spectra of the simulated test set for varying SNRs is illustrated in Figure 4.

**Figure 2:**
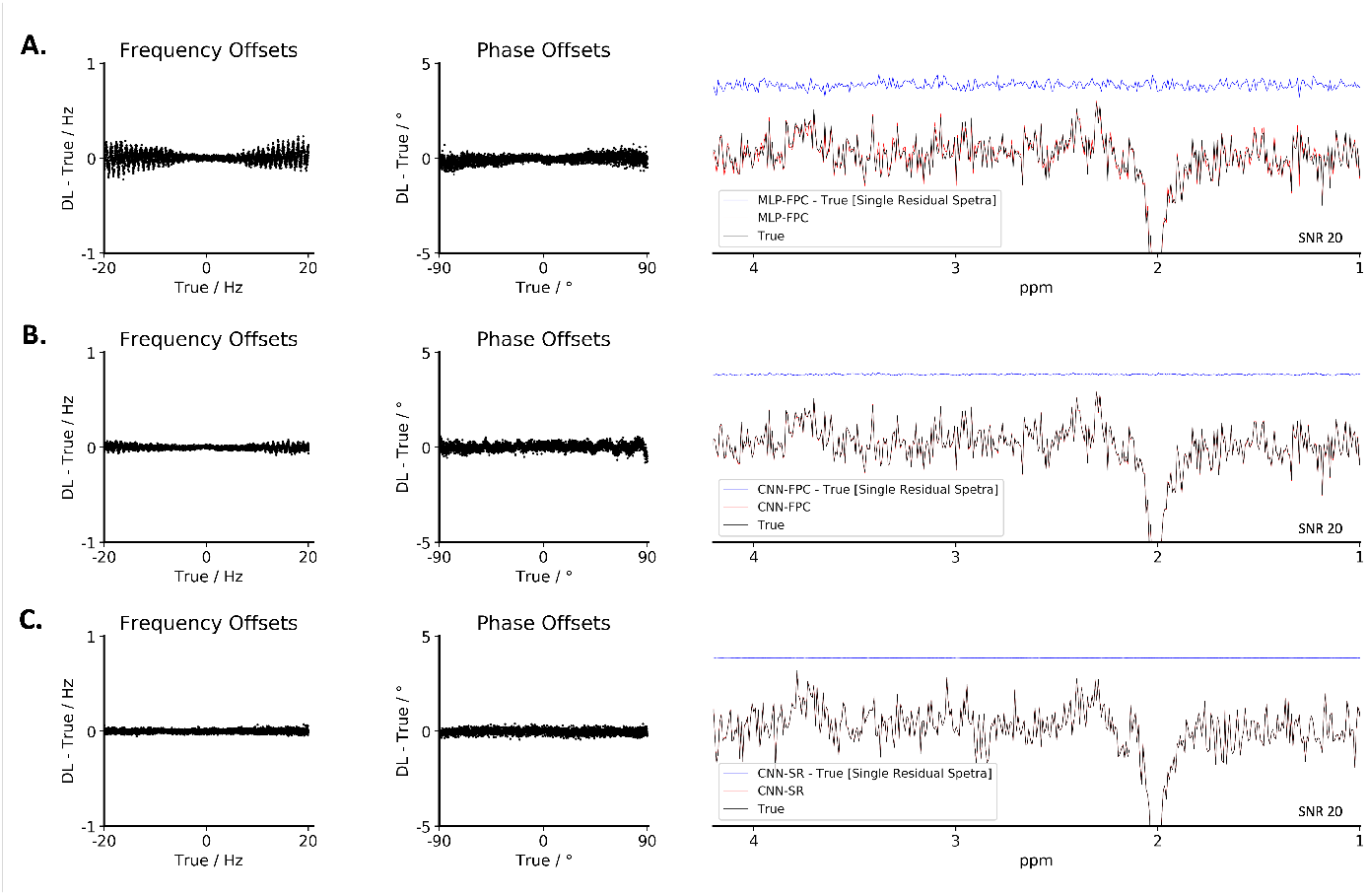
Visualization of the performance of the deep learning models (MLP-FPC, CNN-FPC, CNN-SR) for frequency and phase correction using the published simulated dataset with added noise at the SNR of 20. For the MLP-FPC, the CNN-FPC and the CNN-SR model, the scatter plots on the left show the correction errors between the ground truths and model predictions at different frequency and phase offsets. The spectra on the right demonstrate the spectrum predicted by each deep learning model, the true MEGA-PRESS difference spectra, and the subtraction between them. Among all 3 models, the MLP-FPC exhibits larger correction errors for frequency and phase offset followed by the CNN-FPC, with both being outperformed by the CNN-SR. (A) Output of the MLP-FPC model on the simulated dataset; (B) Output of the CNN-FPC model on the simulated dataset; (C) Output of the CNN-SR model on the simulated dataset.

**Figure 3:**
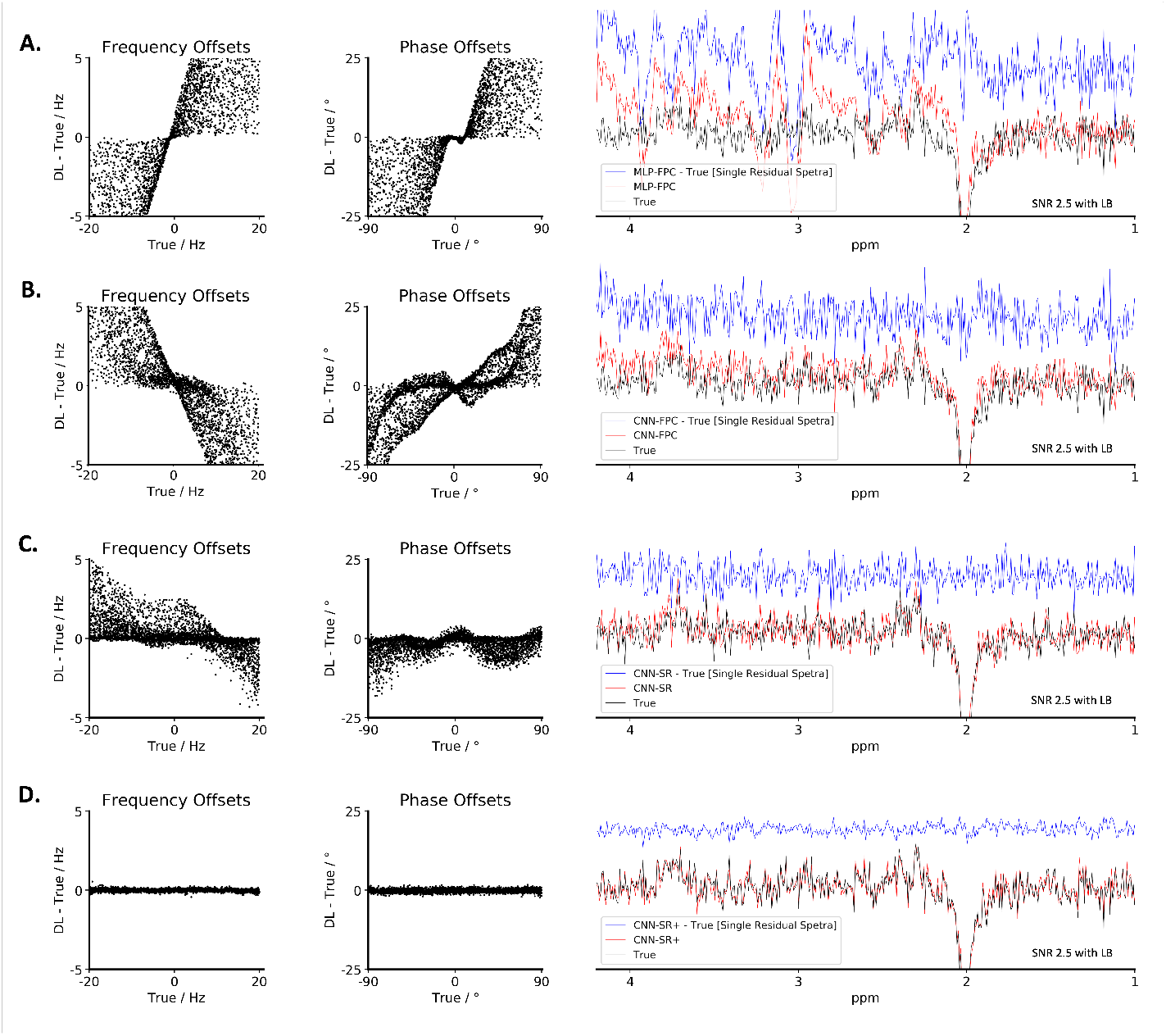
Visualization of the performance of the deep learning models (MLP-FPC, CNN-FPC, CNN-SR, CNN-SR+) for frequency and phase correction using the published simulated dataset with line broadening and added noise at the SNR of 2.5. For the MLP-FPC, the CNN-FPC, the CNN-SR and the CNN-SR+ models, the scatter plots on the left show the correction errors between the ground truths and model predictions at different frequency and phase offsets. The spectra on the right demonstrate the spectrum predicted by each deep learning model, the true MEGA-PRESS difference spectra, and the subtraction between them. Among all 4 models, the MLP-FPC exhibits larger correction errors for frequency and phase offset followed by the CNN-FPC and the CNN-SR, with all being outperformed by the CNN-SR+. (A) Output of the MLP-FPC model on the simulated dataset; (B) Output of the CNN-FPC model on the simulated dataset; (C) Output of the CNN-SR model on the simulated dataset; (D) Output of the CNN-SR+ model on the simulated dataset.

**Figure 4:**
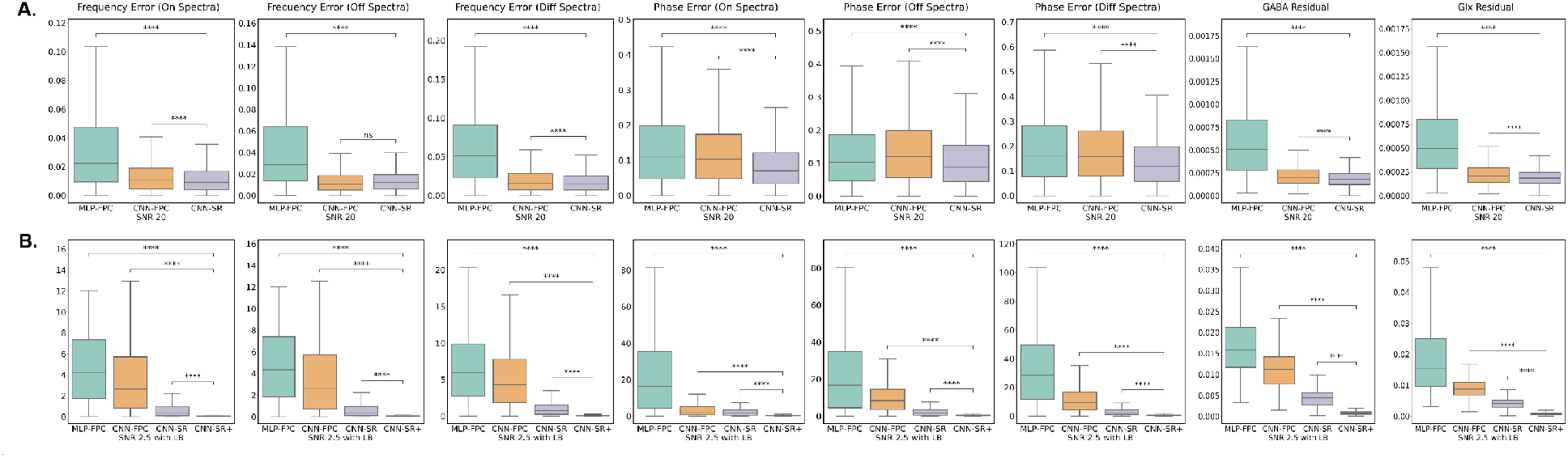
Comparison between the MLP-FPC model, the CNN-FPC model and the CNN-SR model for frequency-and-phase correction of the On, OFF and Diff spectra at the SNR of 20 and the SNR of 2.5 with line broadening. From left to right: the frequency estimation error of the On spectra, the frequency estimation error of the Off spectra, the frequency estimation error of the Diff spectra, the phase estimation error of the On spectra, the phase estimation error of the Off spectra, the phase estimation error of the Diff spectra, the GABA residual spectra mean absolute error and the Glx residual spectra mean absolute error. (A) Box plots showing the frequency estimation error (in Hz), the phase estimation error (in degrees) and the GABA and the Glx residual spectra mean absolute error of the MLP-FPC model, the CNN-FPC model and the CNN-SR model at the SNR of 20; (B) Box plots showing the frequency estimation error (in Hz), the phase estimation error (in degrees) and the GABA and the Glx residual spectra mean absolute error of the MLP-FPC model, the CNN-FPC model and the CNN-SR model at the SNR 2.5 with line broadening. ****: The two-tailed *p*-value is less than 0.0001.

For the test set with SNR 20, the CNN-based approaches showed significantly lower frequency estimation errors than the MLP-based approach, and the CNN-SR model showed the lowest phase estimation errors for the On, Off and Diff spectra (Figure 4A). Taking the Diff spectra as an example, the mean frequency offset error was 0.064 ± 0.052 Hz for the MLP-FPC model, 0.020 ± 0.016 Hz for the CNN-FPC model, and 0.017 ± 0.013 Hz for the CNN-SR model. The mean phase offset error was 0.197 ± 0.159° for the MLP-FPC model, 0.186 ± 0.142° for the CNN-FPC model, and 0.137 ± 0.100° Hz for the CNN-SR model.

With a lower SNR at 2.5 with random 0-20 ms line broadening (Figure 4B), the CNN-SR+ model showed significantly lower frequency and phase estimation errors than the other models for the On, Off and Diff spectra. For example, the mean frequency offset and phase estimation errors for the Diff spectra was 6.658 ± 4.734 Hz, 33.760 ± 26.863° for the MLP-FPC model, 5.264 ± 4.170 Hz, 11.824 ± 9.630° for the CNN-FPC model, 1.067 ± 1.061, 2.987 ± 2.662° Hz for the CNN-SR model and 0.080 ± 0.065, 0.554 ± 0.426° for the CNN-SR+ model.

These results in Figure 2 and Figure 3 show that compared to the MLP-based approaches, the CNN-based approaches had smaller errors within the frequency and phase ranges tested. At the SNR of 20, the CNN-SR model performed better than the MLP-FPC model and the CNN-FPC model. When the SNR decreased to 2.5 and line broadening is applied, the CNN-SR+ model performed better than the MLP-FPC, CNN-FPC and CNN-SR models that had less stable predictions and larger errors.

Additionally, by extracting the spectra interval corresponding to GABA (i.e., 2.8 - 3.2 ppm) and Glx (i.e., 3.55 - 3.95 ppm) from the derived mean difference spectra (Figure 4), these residual spectra errors were found to be lower with the CNN-SR (at the SNR of 20) and CNN-SR+ models (at the SNR of 2.5 with line broadening) than the MLP-FPC and CNN-FPC models. Consequently, the residual spectra errors using CNN-based models for the full spectra were significantly lower than those of the MLP-based model for the On, OFF and Diff spectra at a lower SNR, indicating CNN-based models’ higher performance and robustness in the presence of noise with respect to the MLP-FPC model. Among CNN-based models, the CNN-SR+ model performed the best in terms of frequency and phase estimation errors and noise tolerance, followed by the CNN-SR model. (All numerical results are shown in Supporting Table S1).

### 3.2. Model-performance Evaluation and Spectra Analysis for the in vivo Big GABA Datasets

Figure 5A and Figure 5B illustrate the Off and Diff spectra resulting from the 131 *in vivo* Big GABA Philips datasets without (column 1) or with (columns 2-4) additional artificial offsets for no correction, MLP-FPC model correction, CNN-FPC model correction, and CNN-SR+ model correction. The additional frequency and phase offsets applied to the same 101 datasets are small offsets (0–5 Hz; 0–20°), moderate offsets (5–10 Hz; 20–45°), and large offsets (10–20 Hz; 45–90°). Figure 5C, Figure 5D and Figure 5E demonstrate performance scores comparing the CNN-FPC model to the MLP-FPC model, the CNN-SR+ model to the MLP-FPC model and the CNN-SR+ model to the CNN-FPC model for the 101 *in vivo* datasets.

**Figure 5:**
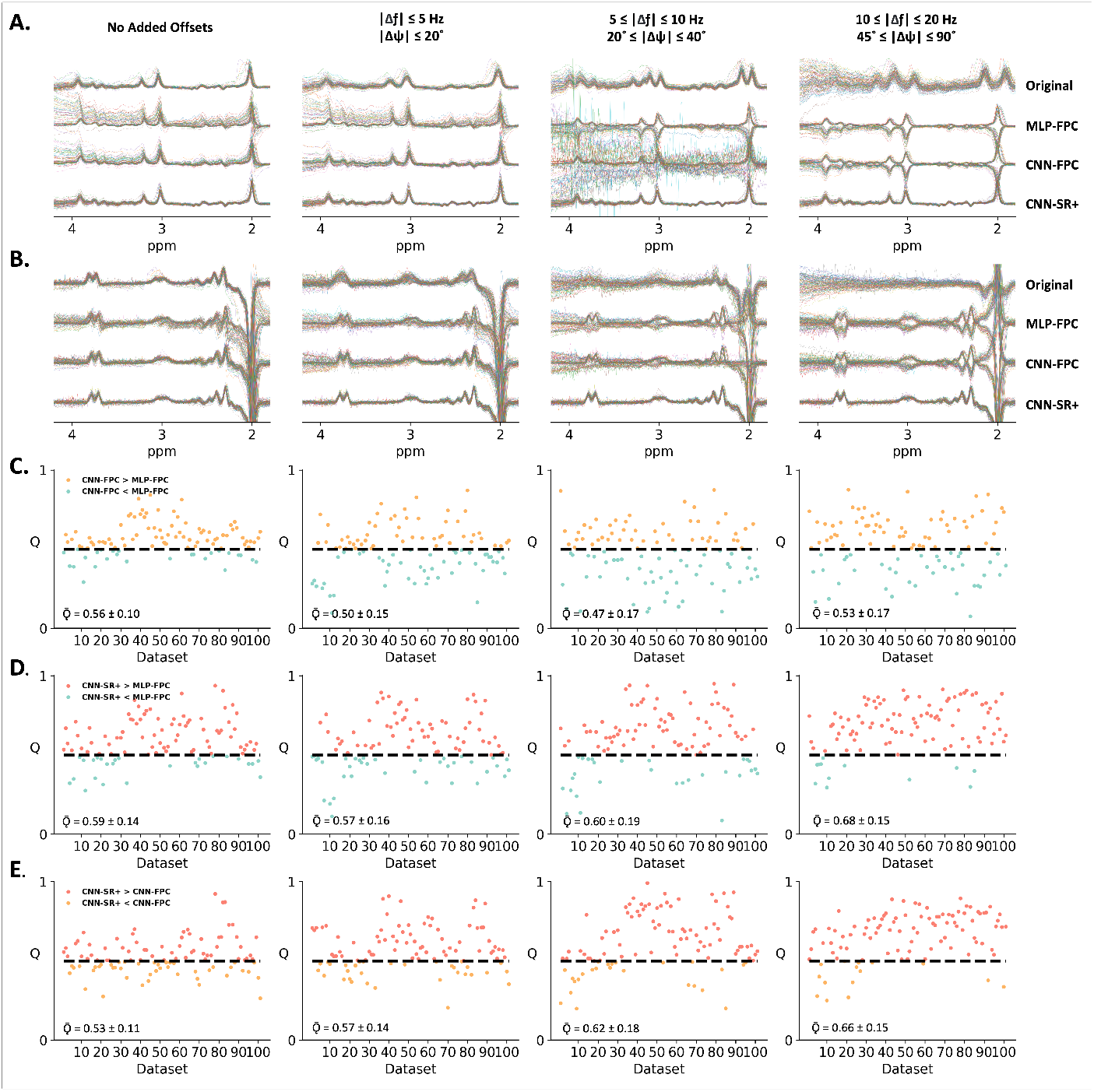
The *in vivo* Off and Diff spectra results of models with different level of added offests and performance scores comparing the CNN-FPC model to the MLP-FPC model, the CNN-SR+ model to the MLP-FPC model and the CNN-SR+ model to the CNN-FPC model for the 101 *in vivo* datasets. (A) The original Off spectra and the results of 3 models after applying corrections to the *in vivo* data without further manipulation and with additional frequency and phase offsets applied to the same 101 datasets: small offsets (0-5 Hz; 0-20°), moderate offsets (5-10 Hz; 20-40°), and large offsets (10-20 Hz; 45-90°); (B) The original Diff spectra and the results of 3 models after applying corrections to the *in vivo* data without further manipulation and with additional frequency and phase offsets applied to the same 101 datasets: small offsets (0-5 Hz; 0-20°), moderat offsets (5-10 Hz; 20-40°), and large offsets (10-20 Hz; 45-90°); (C) Comparative performance Q scores for the CNN-FPC model and the MLP-FPC model for each dataset. A score above 0.5 indicated that the CNN-FPC model performed better than the MLP-FPC model in terms of alignment, whereas a score below 0.5 indicated the opposite. (D) Comparative performance Q scores for the CNN-SR+ model and the MLP-FPC model for each dataset. (E) Comparative performance Q scores for the CNN-SR+ model and the CNN-FPC model for each dataset.

When small offsets were added, the MLP-FPC and CNN-FPC models performed similarly (mean performance score 0.50 ± 0.15, Figure 5C, column2), and both were outperformed by the CNN-SR+ model. The mean performance score of the CNN-SR+ model against the MLP-FPC model was 0.57 ± 0.16 (Figure 5D, column 2) and was 0.57 ± 0.14 for the CNN-SR+ model against the CNN-FPC model (Figure 5E, column 2).

As for moderate offsets, the performance of the MLP-FPC model and the CNN-FPC model was comparable, but the CNN-SR+ model still performed better. The mean performance score of the CNN-FPC model against the MLP-FPC model was 0.47 ± 0.17 (Figure 5C, column 3), while it was 0.60 ± 0.19 for the CNN-SR+ model against the MLP-FPC model (Figure 5D, column 3), and 0.62 ± 0.18 for the CNN-SR+ model against the CNN-FPC model (Figure 5E, column 3).

When large offsets were added, the performance of the CNN-FPC model was slightly better than the MLP-FPC model. The CNN-SR+ model still significantly outperformed the MLP-FPC and CNN-FPC models. The mean performance score of the CNN-FPC model against the MLP-FPC model was 0.53 ± 0.17 (Figure 5C, column 4), while it was 0.68 ± 0.15 for the CNN-SR+ model against the MLP-FPC model (Figure 5D, column 4), and 0.66 ± 0.15 for the CNN-SR+ model against the CNN-FPC model (Figure 5E, column 4).

For small and moderate offsets, the CNN-FPC corrected spectra and MLP-FPC corrected spectra (Figure 5B, columns 2-3) are similar to the original spectra (Figure 5B, column 1). However, for large offsets, the MLP-FPC corrected spectra (Figure 5B, column 4) slightly diverge from the original spectra, while the CNN-FPC corrected spectra still are not noticeably different from the original spectra. Comparably, the CNN-SR+ corrected spectra is always consistent with the original spectra, regardless of the scale of offsets added. The superior performance of the CNN-SR+ model was also indicated by the variances of choline interval for the 101 *in vivo* datasets (Figure 6). With no offset or large offset, the CNN-FPC model had lower choline interval variance than the MLP-FPC model; but with small or medium offsets, the MLP-FPC model had lower choline interval variance than the CNN-FPC model. The CNN-SR+ model, in contrast, had relatively stable performance and its generated variance of the choline interval was significantly lower than both the MLP-FPC model and the CNN-FPC model at all offset levels. Also, the larger the offset, the more significant the CNN-SR+ model was better.

**Figure 6:**
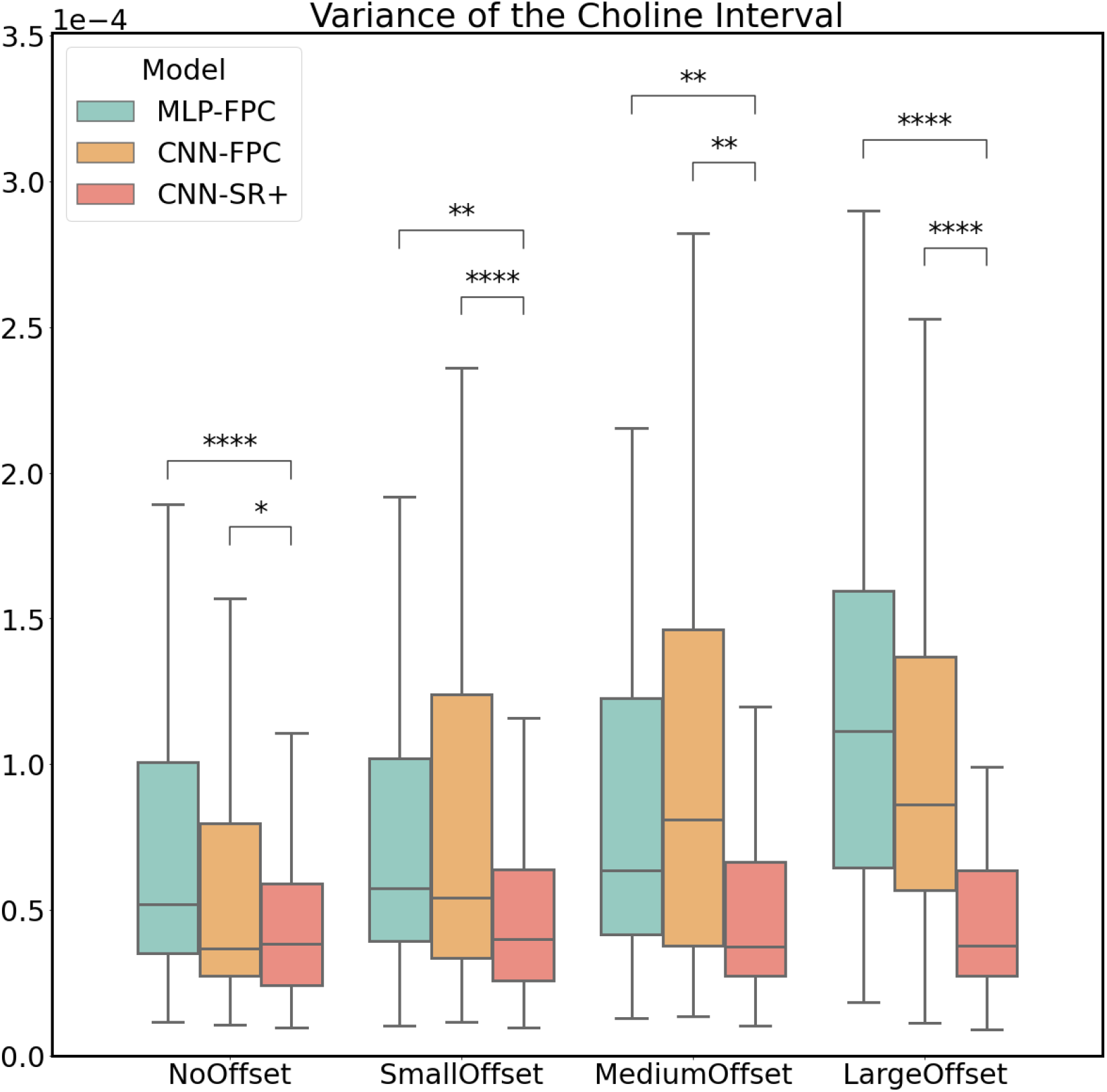
Comparison of the variance of the choline interval in the edited *in vivo* difference spectra among the MLP-FPC, CNN-FPC and CNN-SR+ models with different level of added offsets. From left to right: box plots of choline interval variances with no offset, small offsets, medium offsets and large offsets. The CNN-SR+ model has relatively stable performance and its generated variance of the choline interval is significantly lower than both the MLP-FPC model and the CNN-FPC model at all offset levels. With no offset or large offset, the CNN-FPC model has lower choline interval variance than the MLP-FPC model; but with small or medium offsets, the MLP-FPC model has lower choline interval variance than the CNN-FPC model. ****: The two-tailed *p*-value is less than 0.0001; **: The two-tailed *p*-value is bewteen 0.001 and 0.01; *: The two-tailed *p*-value is between 0.01 and 0.05.

To prove the robustness and spectra quality, experiments were also conducted to compare our best-performed CNN-SR+ model and the published non-deep learning approach, model-based SR (mSR) [2] (Figure 7). mSR exhibited the same performance pattern as our CNN-SR+ model, with a similar mean performance score of 0.50 ± 0.07 for small offsets. They had a similar level of variance of choline interval at around 0.6 *×* 10^*−*4^, with no significant difference.

**Figure 7:**
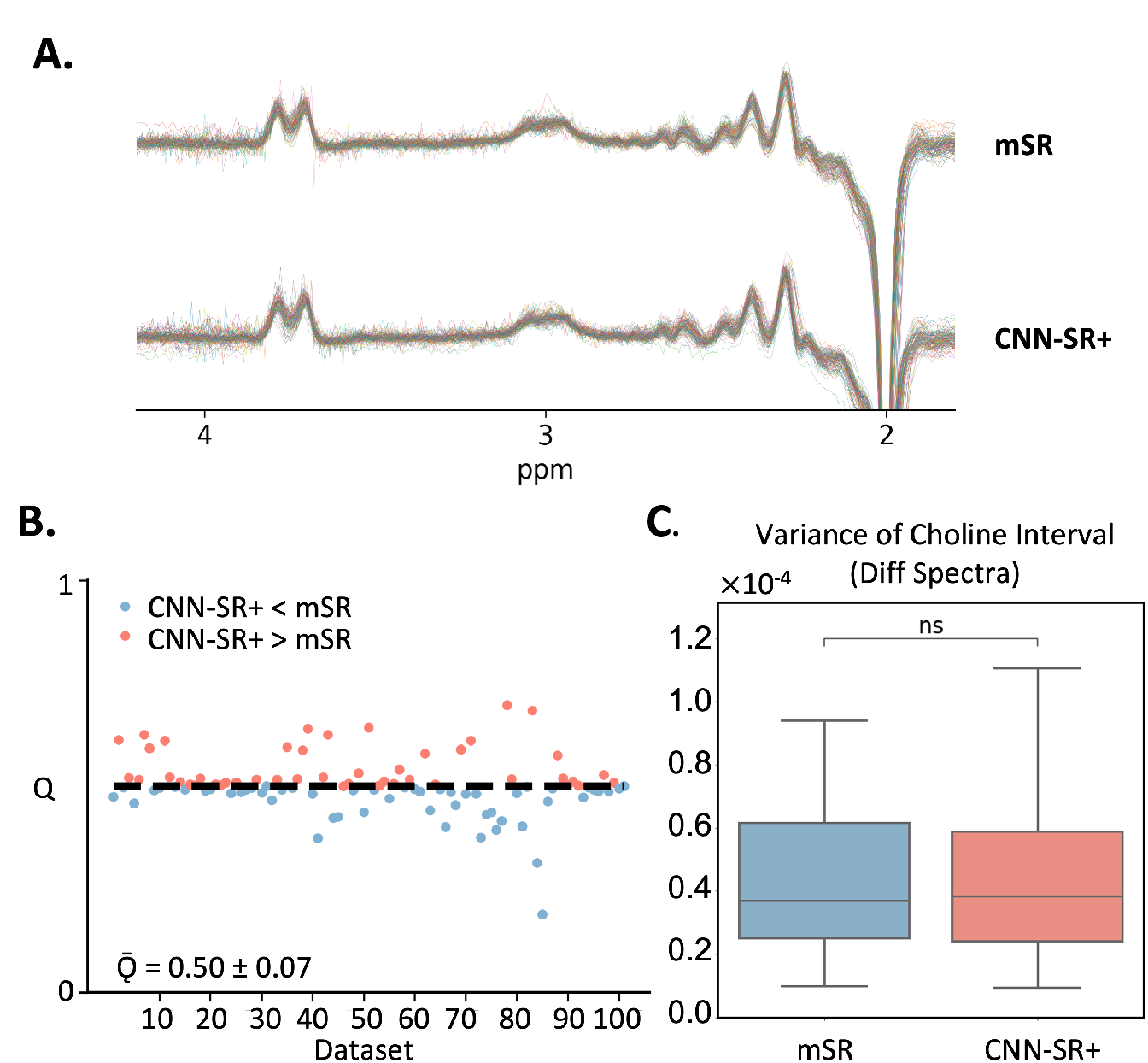
Model performance comparison between the CNN-SR+ model and the mSR model for the *in vivo* datasets, in terms of the Off spectra, Diff spectra, performance scores and the variance of choline interval. (A) The Diff spectra results of the CNN-SR+ model and the mSR model. (B) Comparative performance scores Q for the CNN-SR+ model and the mSR model for each dataset. A score above 0.5 indicated that the CNN-SR+ model performed better than the mSR model in terms of alignment, whereas a score below 0.5 indicated the opposite. (C) Box plots of the variance of the choline interval of the CNN-SR+ model and the mSR model, showing no significant difference.

## 4. Discussion

The metabolic profile of both human and animal brains may be non-invasively and quantitatively measured using MRS. It is extremely beneficial for research and clinical applications since it provides essential information on the metabolic state of the brain. However, the collected data could be affected, since MRS is prone to scanner instability brought on by factors like frequency drift and subject motion. In order to accurately represent and measure metabolites, FPC through spectral registration is a crucial preprocessing step that avoids unwanted spectral distortions which may bias the metabolite quantification.

From Figure 2, 3 and 4, the results show that our proposed CNN-SR model is more robust and has a better performance when compared to the other sequential FPC deep learning methods (CNN-FPC and MLP-FPC) for all testing conditions in the simulated data. Figure 2 and 4 demonstrate that in simulated data with SNR 20, the MLP-FPC model exhibits larger correction errors for frequency and phase offset, and On/OFF mismatch errors followed by the CNN-FPC model, with both being outperformed by the CNN-SR model. The scatter plots also demonstrate that CNN-SR model offset predictions are more congregated near the y=0, indicating better prediction accuracy than the FPC models. The single residual spectra that plots the difference between the prediction and the ground truth also exhibits consistent results, where a more complete subtraction (straight line) is obtained. Likewise, Figure 3 and 4 confirm that the CNN-SR model surpasses both sequential FPC deep learning methods when faced with more distorted data (SNR 2.5 with LB 0-20ms). The results show that CNN-SR had smaller mean absolute errors for both frequency and phase offsets predictions and Diff spectra derivation evidencing this approach is more robust to noise and provides more accurate predictions.

Moreover, the performance of CNN-SR can be significantly improved by fine-tuning the model to the specific data. Given the nature of *in vivo* data where ground truths are not present and the data is disturbed by multiple variables (i.e. noise, subject motion), an unsupervised learning spectral registration approach is considered to further enhance the model performance. A more refined model (CNN-SR+) was further trained with the more distorted data from the pre-trained the CNN-SR model. In Figure 3 and 4, it is clear how the CNN-SR+ model significantly improves the results, as seen in the smaller correction errors for both phase and frequency, and in the subtraction of the prediction to the ground truth Diff spectra being a straight line.

When testing on *in vivo* data for different phase and frequency offsets, the utility of an unsupervised spectral registration model, the CNN-SR+ model demonstrated once again to have superior performance. Figure 5 shows that OFF, and Diff spectra are clearer for this model across all testing conditions, with shapes and peaks better preserved. Furthermore, Q scores are consistently higher using the CNN-SR+ model in comparison to all the other FPC deep learning models. Nevertheless, these results remain equivalent to mSR, the state of art non-deep learning numerical correction method, as shown in Supplementary Figure 1.

These findings illustrate the value of deep learning for spectral registration and evince the utility and strengths of simultaneous FPC with a spectral registration model framework. The CNN-SR model can perform simultaneous frequency and phase correction, and compared to the CNN-FPC and MLP-FPC models, it produces more reliable, robust and accurate results in a shorter processing time and with higher computational efficiency. Additionally, our framework has the remarkable capability of adopting an unsupervised learning approach. Contrary to other FPC models that would need a ground truth, this approach can take advantage of using spectra loss to learn in an unsupervised manner. This is a powerful advantage, widely applicable to training and testing on *in vivo* data. Additionally, from Supplementary Figure 1, it can also be shown that the performance of CNN-SR+ is comparable to mSR, but given advantages as stated previously (shorter processing time and higher computational efficiency), our model surpasses the utility of mSR. Results in this study reveal that by employing unsupervised learning we were able to fine-tune our model to obtain state of the art performance for any given dataset.

Despite the utility of our framework, there still remain several limitations and lines of research to further explore. This study was solely conducted on humans, but MRS is a widely available approach for animals as well, playing a noteworthy role in pre-clinical studies. Further exploration on animal data could be carried out in the future to validate the generalizability of our framework. Additionally, testing was conducted on *in vivo* data, unfolding the possibility to also consider other living conditions like *in situ, ex vivo* and *in vitro*. Regarding the JDE sequences, our model was tested on MEGA-PRESS, but other sequences such as PRESS and MEGA-sLASER could be considered in the future. Furthermore, data from vendors like GE and Siemens is publicly available, so it would be valuable to work with more vendor datasets. Similarly, different magnetic field strengths other than 3T (e.g. 9T, 12T) are important variables to take into account in the future. In this study we only considered frequency and zero-order phase. However, inclusion of other parameters like first-order phase, amplitude and bandwidth variance in different transients could be examined in upcoming research studies. Although CNN-SR outperformed the other deeplearning approaches, it is still comparable to the state of the art model (mSR). However, given the advantages our deep learning approach provides when conducting spectral registration, such as its high computational efficiency, quick processing time and being able to generalize well on datasets from different modalities, our approach has more utility for the users. Our model can also adopt to very small datasets and can be applied to the same dataset numerous times demonstrating overcoming potential issues such as lack of resources. For future work, creating a model that bests mSR should be explored. A way to approach this could include contemplating other backbone frameworks, such as transformers. Finally, it would be of great interest to demonstrate our model’s clinical utility by processing MRS spectra from patients with neuropsychiatric disorders.

## 5. Conclusion

This work provides the first proof of concept of the feasibility of a CNN framework for MRS spectra registration with both supervised and unsupervised learning. Our CNN-SR model shows better performance and delivers results more robust to noise as compared to the current state-of-the-art models in both simulation and *in vivo* tests.

## Data and Code Availability Statement

The CNN-SR code used in this project is proprietary. The CNN code is available at https://github.com/davidjma/CNN-SR. The CNN code is © 2022 The Trustees of Columbia University in the City of New York. This work may be reproduced and distributed for academic non-commercial purposes only.

## Acknowledgments

This study was performed at the Zuckerman Mind Brain Behavior Institute MRI Platform, a shared resource. The results published here are based on public Big GABA data obtained from 24 research sites. The authors appreciate Nuriel Tal for his constructive comments and suggestions.

## Figures and Tables

**Table 1:** Table of Models’ Performance for the Simulated Datasets. Table of mean absolute errors of the MLP-FPC, CNN-FPC, CNN-SR and CNN-SR+ model for frequency correction, phase correction, GABA residal and Glx residual on the simulation dataset with different level of noise

| Simulation Dataset | Model | Frequency Error (Hz) |  |  | Phase Error (Degree) |  |  | GABA Residual | Glx Residual |
| --- | --- | --- | --- | --- | --- | --- | --- | --- | --- |
|  |  | On | Off | On/Off Mismatch | On | Off | On/Off Mismatch |  |  |
| SNR 20 | MLP-FPC | 0.034 ± 0.033 | 0.043 ± 0.039 | 0.064 ± 0.052 | 0.140 ± 0.119 | 0.132 ± 0.116 | 0.197 ± 0.159 | 0.00059 ± 0.00039 | 0.00058 ± 0.00036 |
|  | CNN-FPC | 0.014 ± 0.012 | 0.014 ± 0.012 | 0.020 ± 0.016 | 0.125 ± 0.104 | 0.141 ± 0.106 | 0.186 ± 0.142 | 0.00022 ± 0.00012 | 0.00023 ± 0.00012 |
|  | CNN-SR | 0.011 ± 0.009 | 0.014 ± 0.010 | 0.017 ± 0.013 | 0.085 ± 0.067 | 0.104 ± 0.076 | 0.137 ± 0.100 | 0.00020 ± 0.00010 | 0.00019 ± 0.00009 |
| SNR 2.5<br>with Random 0-20 ms LB | MLP-FPC | 4.643 ± 3.204 | 4.715 ± 3.221 | 6.658 ± 4.734 | 22.247 ± 20.680 | 22.063 ± 20.122 | 33.760 ± 26.863 | 0.01694 ± 0.00729 | 0.01827 ± 0.01066 |
|  | CNN-FPC | 3.599 ± 3.225 | 3.465 ± 3.126 | 5.264 ± 4.170 | 4.155 ± 5.389 | 10.468 ± 8.931 | 11.824 ± 9.630 | 0.01103 ± 0.00400 | 0.00938 ± 0.00360 |
|  | CNN-SR | 0.670 ± 0.867 | 0.678 ± 0.883 | 1.067 ± 1.061 | 2.297 ± 2.429 | 2.367 ± 2.616 | 2.987 ± 2.662 | 0.00438 ± 0.00214 | 0.00404 ± 0.00174 |
|  | CNN-SR+ | 0.053 ± 0.044 | 0.058 ± 0.050 | 0.080 ± 0.065 | 0.366 ± 0.293 | 0.416 ± 0.317 | 0.554 ± 0.426 | 0.00084 ± 0.00046 | 0.00085 ± 0.00044 |

**Table 2:** Table of Q Scores for the *in vivo* Datasets. Table of performance scores Q calculated between MLP-FPC, CNN-FPC and CNN-SR+ model under four conditions: no added offsets, small offsets (C1), moderat offsets (C2) and large offsets (C3).

| Model | Q Score |  |  |  |
| --- | --- | --- | --- | --- |
|  | No Added Offsets | C1 | C2 | C3 |
| CNN-FPC vs. MLP-FPC | 0.56 ± 0.10 | 0.50 ± 0.15 | 0.47 ± 0.17 | 0.53 ± 0.17 |
| CNN-SR+ vs. MLP-FPC | 0.59 ± 0.14 | 0.57 ± 0.16 | 0.60 ± 0.19 | 0.68 ± 0.15 |
| CNN-SR+ vs. CNN-FPC | 0.53 ± 0.11 | 0.57 ± 0.14 | 0.62 ± 0.18 | 0.66 ± 0.15 |

**Table 3:** Table of Choline Residuals for the *in vivo* Datasets. Table of choline residuals calculated on the MLP-FPC, CNN-FPC and CNN-SR+ model under four conditions: no added offsets, small offsets (C1), moderat offsets (C2) and large offsets (C3).

| Model | Variance of the Choline Interval |  |  |  |
| --- | --- | --- | --- | --- |
|  | No Added Offsets | C1 | C2 | C3 |
| MLP-FPC | 0.000086 ± 0.000084 | 0.000094 ± 0.000088 | 0.000029 ± 0.000025 | 0.000126 ± 0.000089 |
| CNN-FPC | 0.000074 ± 0.000103 | 0.000129 ± 0.000198 | 0.000049 ± 0.000100 | 0.000132 ± 0.000188 |
| CNN-SR+ | 0.000055 ± 0.000054 | 0.000062 ± 0.000068 | 0.000033 ± 0.000023 | 0.000067 ± 0.000102 |

## References

1 Liu C, Ma D. J., et al (2021) JET - A Matlab toolkit for automated J-difference-edited MR spectra processing of in vivo mouse MEGA-PRESS study at 9.4T. In Joint Annual Meeting ISMRM & SMRT 2021 (Vancouver, Canada).

2 Near J, Edden R, Evans CJ, Paquin R, Harris A, Jezzard P. Frequency and phase drift correction of magnetic resonance spectroscopy data by spectral registration in the time domain. Magn Reson Med. 2015;73:44–50.

3 Oeltzschner G, Zöllner HJ, Hui SCN, et al. Open-source processing, reconstruction and estimation of magnetic resonance spectroscopy data. J Neurosci Methods. 2020;343:108827.

4 Rothman DL, Petroff OA, Behar KL, Mattson RH. Localized 1H NMR measurements of gamma-aminobutyric acid in human brain in vivo. Proc Natl Acad Sci USA 1993;90(12):5662–5666.

5 Mescher M, Merkle H, Kirsch J, Garwood M, Gruetter R. Simultaneous in vivo spectral editing and water suppression. NMR Biomed 1998;11(6):266–72.

6 Mullins PG, McGonigle DJ, O’Gorman RL, et al. Current practice in the use of MEGA-PRESS spectroscopy for the detection of GABA. Neuroimage 2014;86:43–52.

7 Sawiak SJ, Jupp B, Taylor T, Caprioli D, Carpenter TA, Dalley JW. in vivo γ-aminobutyric acid measurement in rats with spectral editing at 4.7 T. J Magn Reson Imaging 2016;43(6):1308–1312.

8 Fu Y, Lei Y, Wang T, Higgins K, Bradley JD, Curran WJ, Liu T, Yang X. LungRegNet: An unsupervised deformable image registration method for 4D-CT lung. Med Phys. 2020 Apr;47(4):1763–1774. doi: 10.1002/mp.14065. Epub 2020 Feb 26. PMID: 32017141; PMCID: PMC7165051.

9 Chen, J., Frey, E.C., Du, Y. (2022). Unsupervised Learning of Diffeomorphic Image Registration via TransMorph. In: Hering, A., Schnabel, J., Zhang, M., Ferrante, E., Heinrich, M., Rueckert, D. (eds) Biomedical Image Registration. WBIR 2022. Lecture Notes in Computer Science, vol 13386. Springer, Cham.

10 Tapper S, Mikkelsen M, Dewey BE, Zöllner HJ, Hui SCN, Oeltzschner G, Edden RAE. Frequency and phase correction of J-difference edited MR spectra using deep learning. Magn Reson Med. 2021 Apr;85(4):1755–1765. doi: 10.1002/mrm.28525. Epub 2020 Nov 18. PMID: 33210342.

11 Ma DJ, L. Ha, Ye Y, Laine AF, Lieberman JA, Rothman DL, Small SA, Guo J. MR spectroscopy frequency and phase correction using convolutional neural networks. Magn Reson Med. 2022 Apr;87(4):1700–1710. doi: 10.1002/mrm.29103. Epub 2021 Dec 21. PMID: 34931715.

12 Mikkelsen M, Barker PB, Bhattacharyya PK, et al. Big GABA: Edited MR spectroscopy at 24 research sites. Neuroimage. 2017;159:32–45.

13 D. Kingma and J. Ba. Adam: A method for stochastic optimization”. arXiv 2014, arXiv:1412.6980.

